# Developmental stage- and site-specific transitions in lineage specification and gene regulatory networks in human hematopoietic stem and progenitor cells

**DOI:** 10.1101/2021.04.20.440420

**Authors:** Anindita Roy, Guanlin Wang, Deena Iskander, Sorcha O’Byrne, Natalina Elliott, Jennifer O’Sullivan, Gemma Buck, Elisabeth F. Heuston, Wei Xiong Wen, Alba Rodriguez Meira, Peng Hua, Anastasios Karadimitiris, Adam J Mead, David Bodine, Irene Roberts, Bethan Psaila, Supat Thongjuea

## Abstract

Human hematopoiesis is a dynamic process that starts *in utero* 4 weeks post-conception. Understanding the site- and stage-specific variation in hematopoiesis is important if we are to understand the origin of hematological disorders, many of which occur at specific points in the human lifespan. To unravel how the hematopoietic stem/progenitor cell (HSPC) compartments change during human ontogeny and the underlying gene regulatory mechanisms, we compared 57,489 HSPCs from 5 different tissues spanning 4 developmental stages through the human lifetime. Single-cell transcriptomic analysis identified significant site- and developmental stage-specific transitions in cellular architecture and gene regulatory networks. Uncommitted stem cells showed progression from cycling to quiescence and increased inflammatory signalling during ontogeny. We demonstrate the utility of this dataset for understanding aberrant hematopoiesis through comparison to two cancers that present at distinct timepoints in postnatal life – juvenile myelomonocytic leukemia, a childhood cancer, and myelofibrosis, which classically presents in older adults.

## INTRODUCTION

In the human embryo, definitive blood stem cells first arise in the aorta-gonad-mesonephros (AGM) region at 4-5 weeks post conception. Hematopoiesis then migrates to the fetal liver (FL) and subsequently to the bone marrow (BM), which becomes the dominant hematopoietic organ at birth and remains so throughout postnatal life (Ivanovs et al., 2017). Single cell transcriptomics have been extensively applied to clarify the cellular architecture and molecular pathways in hematopoiesis, but the majority of studies have been done in adult tissues (Hay et al., 2018; Velten et al., 2017), mouse models (Dahlin et al., 2018; Tusi et al., 2018) or human cord blood (Notta et al., 2016; Zheng et al., 2018). A recent analysis of 1^st^ and 2^nd^ trimester human fetal liver (FL), fetal kidney and fetal skin indicated that the hematopoietic compartment in FL changes from being predominantly erythroid in early gestation, to lympho-myeloid in later development, and that hematopoietic stem/progenitor cells (HSPCs) become less proliferative during fetal maturation (Popescu et al., 2019). Transition of HSPCs from proliferative to quiescent state has also been associated with migration of hematopoiesis from FL to fetal bone marrow (FBM), along with a concomitant decrease in the frequency of non-committed HSPCs, suggesting a role for the niche in regulating hematopoietic cell state (Ranzoni et al., 2021).

However, the transcriptome profiles of the lineage-negative (Lin-), CD34+ HSPC compartments in hematopoietic tissues during development and postnatal ageing have never been directly compared using the same platform. These studies are crucial if we are to understand the variation in hematopoiesis during normal human ontogeny. For example, whether certain patterns of fetal, pediatric or adult-specific gene expression signatures exist that are permissive for the emergence of age-dependent hematopathologies has not previously been described.

We therefore generated a comprehensive dataset encompassing 57,489 HSPCs sampled from healthy human hematopoietic tissues including first trimester early FL (eFL), paired second trimester FBM and FL (isolated from the same fetuses), pediatric BM (PBM), and adult BM (ABM). To our knowledge, this is the first study that directly compares human HSPC from all stages of ontogeny (early fetal life to adulthood) at single cell level. Precise delineation of the consistencies and differences in the cellular composition and molecular pathways in the HSPC compartments across human development demonstrated pronounced site- and developmental stage-specific transitions in cellular architecture and transcriptional profiles between hematopoietic tissues. While megakaryo-erythropoiesis predominated in early fetal liver, lympho-myeloid progenitors showed dramatic expansion following onset of hematopoiesis in the BM of the developing fetus. The proportion of lymphoid progenitors in the BM hematopoietic compartment then progressively decreased during postnatal life. In addition, HSCs but not lineage-committed progenitors showed progression from cycling to quiescence and increased inflammatory signalling during development from fetal through to adult life. Finally, comparison of HSPCs sampled over normal human ontogeny to HSPCs from two hematological cancers that occur at extremes of life, Juvenile Myelomonocytic Leukaemia (JMML) affecting young children and myelofibrosis (MF) affecting older adults, suggested a fetal origin for JMML and exacerbation of the adult BM-associated inflammatory signalling programs in myelofibrosis HSPCs. These observations highlight the value of this comprehensive single cell transcriptomic resource for understanding the changes in normal hematopoiesis through human ontogeny, as well as unravelling the cellular and molecular disruptions that occur in hematopoietic disorders.

## RESULTS

### Analysis of 57,489 HSPCs revealed 21 distinct cell clusters across human ontogeny

Single-cell RNA sequencing (scRNAseq) was performed on Lin-CD34+ cells from eFL (15,036 cells), matched FL (17,351 cells) and FBM (14,935 cells) isolated from the same fetuses, PBM (13,311 cells) and ABM (6,300 cells) using the 10x Genomics platform (Fig.1A). 66,933 HSPC were captured, and following quality control 57,489 single-cell transcriptomes were included in downstream analyses (Suppl. Table 1). Datasets were integrated using an adaptation of the Harmony algorithm (Korsunsky et al., 2019) that automatically groups marker genes for each cluster per sample, and builds up a gene set database for GSEA analysis (see methods) as implemented in the SingCellaR package (https://github.com/supatt-lab/SingCellaR). This led to optimal data integration with improved resolution of erythroid from eosinophil/basophil/mast progenitors than when using standard Harmony or Seurat methods as indicated by visual inspection (Suppl. Fig. 1A-C) and the objective measures kBET, iLISI and cLISI (Suppl. Fig. 1D-E) (Buttner et al., 2019; Korsunsky et al., 2019).

**Figure 1.**
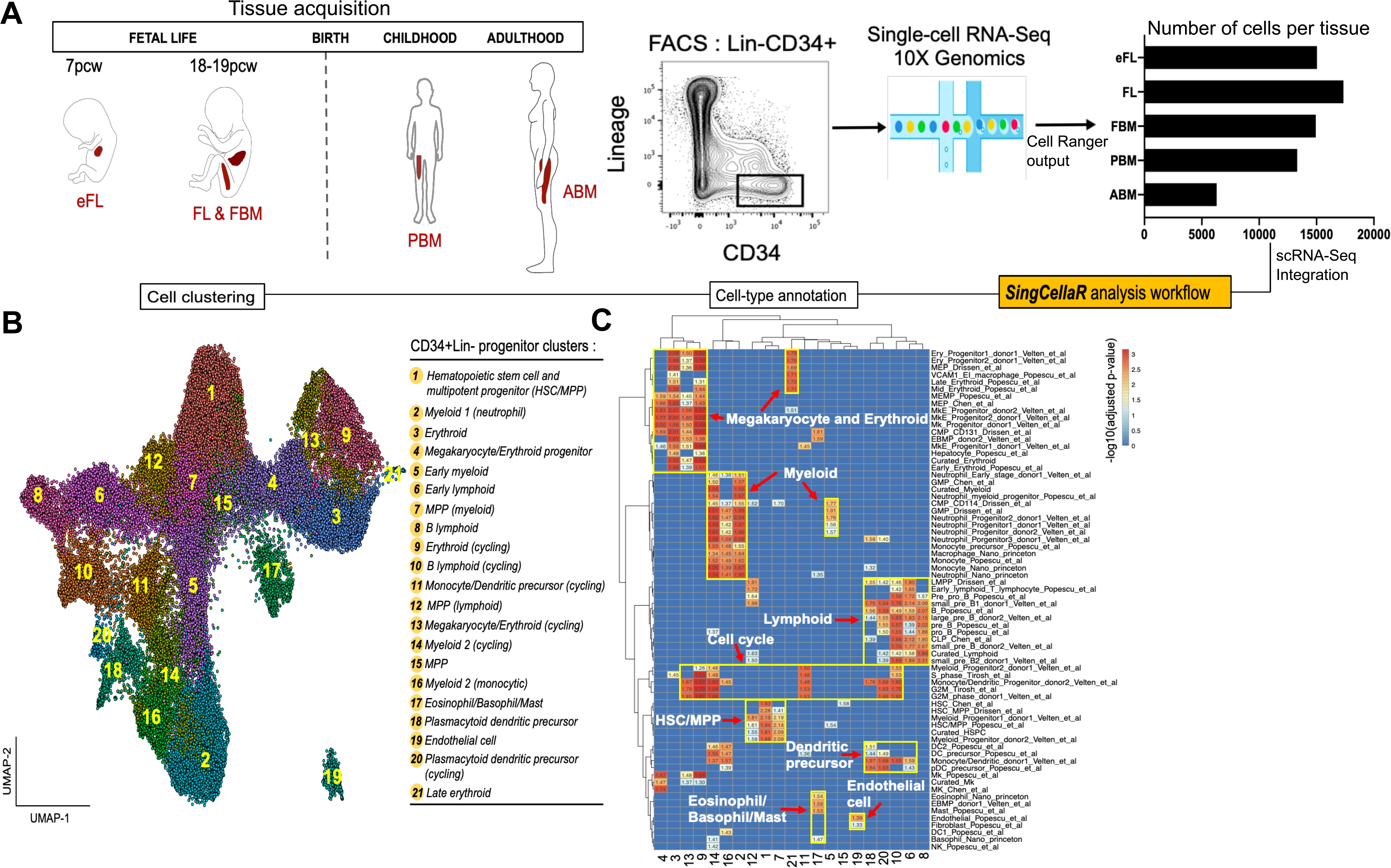
Single cell RNA-sequencing of 57,489 hematopoietic stem/progenitor cells (HSPCs) from five hematopoietic tissues across human ontogeny reveals 21 distinct cellular subsets. **(A)** Experimental design. Lineage-negative (Lin-) CD34+ HSPCs were FACS-sorted for single-cell RNA-sequencing from first trimester fetal liver (eFL), matched second trimester FL and fetal bone marrow (FBM), pediatric bone marrow (PBM), and adult bone marrow (ABM). A bar chart shows the number of cells per tissue. **(B)** Louvain community-detection clustering based on the weighted graph network Uniform Manifold Approximation and Projection (UMAP) of 57,489 cells identified 21 distinct hematopoietic progenitor populations. **(C)** Heatmap generated using SingCellaR’s cell-type annotation system showing positive gene set enrichment scores for each cluster, facilitates cluster identification. X-axis represents clusters as numbered in Fig. 1B. Y-axis represents a curated list of hematopoietic lineage-specific gene sets (Suppl. Table 2)

To identify the cellular composition of the 57,489 HSPCs captured from different stages and tissues over ontogeny, graph-based Louvain clustering was performed. This identified 21 clusters with distinct expression patterns of canonical stem/progenitor and hematopoietic lineage marker genes (Fig. 1B, Suppl. Fig. 2A-B and Suppl. Table 2) (Drissen et al., 2019; Hay et al., 2018; Pellin et al., 2019; Popescu et al., 2019; Velten et al., 2017). While some clusters were easy to identify by their clear expression of canonical markers, other clusters were less readily classifiable due to low-level expression of lineage-affiliated genes and/or expression of multiple lineage markers. To facilitate annotation of these clusters, we collated lineage signature genes from 75 published human hematopoiesis gene sets (Supp. Table 3), and gene set enrichment analyses (GSEA) were performed on the expressed genes for each cluster using these gene sets to guide assignment of cell types (Fig. 1C).

Cluster 1 had robust expression of hematopoietic stem cell/multipotent progenitor (HSC/MPP)-affiliated genes, while clusters 7 and 12 showed HSC/MPP-genes together with evidence of early myeloid or lymphoid priming respectively (Fig. 1B, 1C and Suppl. Fig. 2C). Cell clusters representing the major hematopoietic lineage progenitor subsets (erythroid, myeloid, lymphoid, megakaryocytic, dendritic, eosinophil/basophils/mast cells) were identified, as well as clusters with expression of genes representing >1 lineage likely to represent uncommitted/oligopotent progenitors (e.g., erythro-megakaryocytic [cluster 4] and eosinophil/basophil/mast cell precursors [cluster 17]) (Fig. 1C). A population of non-hematopoietic cells (Cluster 19) expressing endothelial genes was detected, which derived predominantly but not exclusively from eFL. Some lineage progenitor clusters had differential enrichment of G2M checkpoint and S-phase gene signatures (Suppl Table 4), allowing us to classify certain progenitor populations by their distinct proliferation state (e.g. cluster 9 – erythroid (cycling); cluster 10 – B lymphoid (cycling); Fig. 1B).

### Site- and developmental stage-specific differences in the cellular composition of the HSPC compartment

To accurately determine differences in the composition of Lin-CD34+ progenitors between hematopoietic sites and ontological stages, we identified cells showing enriched expression of curated gene signatures (Suppl. Table 4) corresponding to uncommitted HSC/MPP and to the main lineage progenitor sub-types (myeloid, mega-erythroid-mast cell and B-lymphoid progenitors, Fig. 2A). Using an AUCell (Aibar et al., 2017) score of > 0.15, clear distinction between cellular subsets was seen, with 24% of cells (13,871 cells) classified as uncommitted HSC/MPP, 15% as myeloid (8,330 cells), 16% as lymphoid (9,473 cells), 2% as eosinophil/basophil/mast cell (1,361 cells), 12% as erythroid (6,778 cells), and 1% as megakaryocytic (572 cells) progenitors (Fig. 2A).

**Figure 2.**
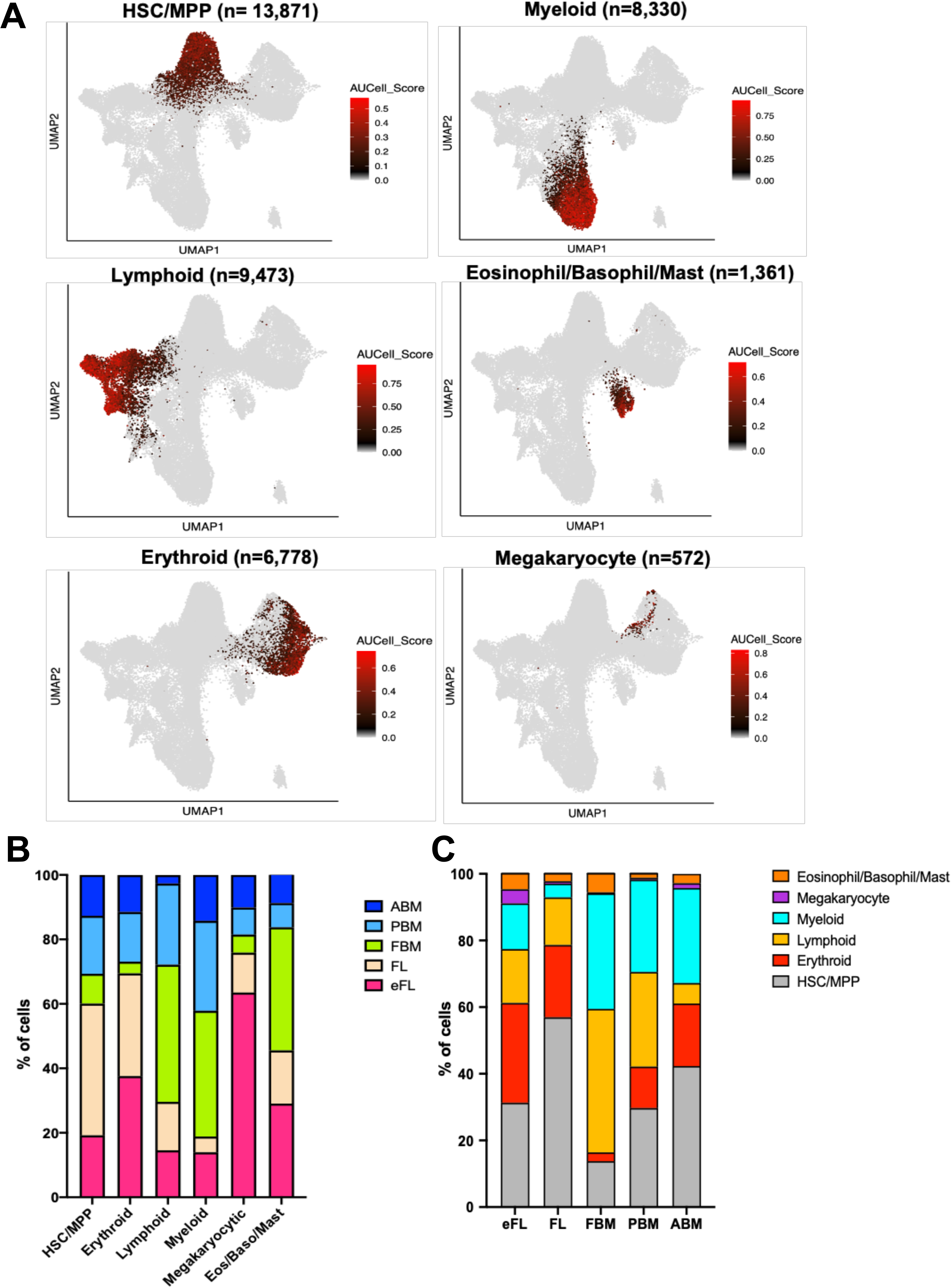
Site and developmental stage-specific changes in composition of the HSPC compartment. **(A)** Cells were classified using an AUCell score of > 0.15 for canonical marker genes for HSC/MPP, myeloid, lymphoid, eosinophil/basophil/mast cell, erythroid, and megakaryocyte progenitors. **(B)** The proportion of HSC/MPP and lineage progenitor subsets that derived from each tissue type. **(C)** The proportion of HSPCs from each tissue classified as HSC/MPP or lineage progenitors subtypes.

1^st^ and 2^nd^ trimester FL contributed the majority (69.4%) of the erythroid progenitors captured (Fig. 2B), whereas lymphoid and myeloid progenitors predominantly derived from BM samples (70.4% and 81.2% respectively, Fig. 2B). Marked differences were observed in origin of the megakaryocyte progenitor cells captured, with a majority (63.5%) captured from eFL (Fig. 2B), while only 5%-12% derived from any other individual tissue. Differences in lineage specification were also demonstrable when the proportions of progenitor cell types were quantified within individual tissue (Fig. 2C and Suppl. Fig. 3A). These differences in lineage specification for the main lineage subtypes described above were maintained after adjusting for the variable cell numbers obtained from each tissue type (Suppl. Fig. 3B). Of note, marked differences were seen in the cellular composition of matched FL and FBM HPSCs when analysing cells representing uncommitted HSPC and the main lineage subtypes. There were 4.4-fold more uncommitted progenitors and 8.7-fold more erythroid progenitors in FL than FBM. The opposite trend was true for myeloid and lymphoid progenitors, which were 7.9-fold and 2.8-fold more frequent in FBM than FL (Fig. 2C).

### Four main differentiation trajectories are present among HSPCs across human ontogeny

To explore the developmental relationships between cell clusters, cells were ordered based on their gene expression using a force directed graph (FDG) network (Fig. 3A – 3C). ‘Lineage signature gene scores’ were defined as before (Psaila et al., 2020) and superimposed on the FDG. This demonstrated four major paths emerging from the HSC/MPP cluster (cluster 1, Fig. 3A), representing erythroid, megakaryocytic, lymphoid, and myeloid differentiation trajectories along pseudotime (Fig. 3B and C). As expected, clusters with multipotent potential (Clusters 1, 5, 7, 11, 12, and 15) were located at the apex or central positions in the trajectories (Fig. 3A and B, grey cells). The existence of the main trajectories was confirmed using diffusion mapping as an alternative trajectory analysis (Suppl. Fig. 3C). Dendritic cell precursors (clusters 18 and 20) were closely affiliated with the lymphoid trajectory, while the trajectory of eosinophil/mast cell/basophil progenitors (cluster 17) was associated with that of erythroid progenitors (Fig. 3A and B).

**Figure 3.**
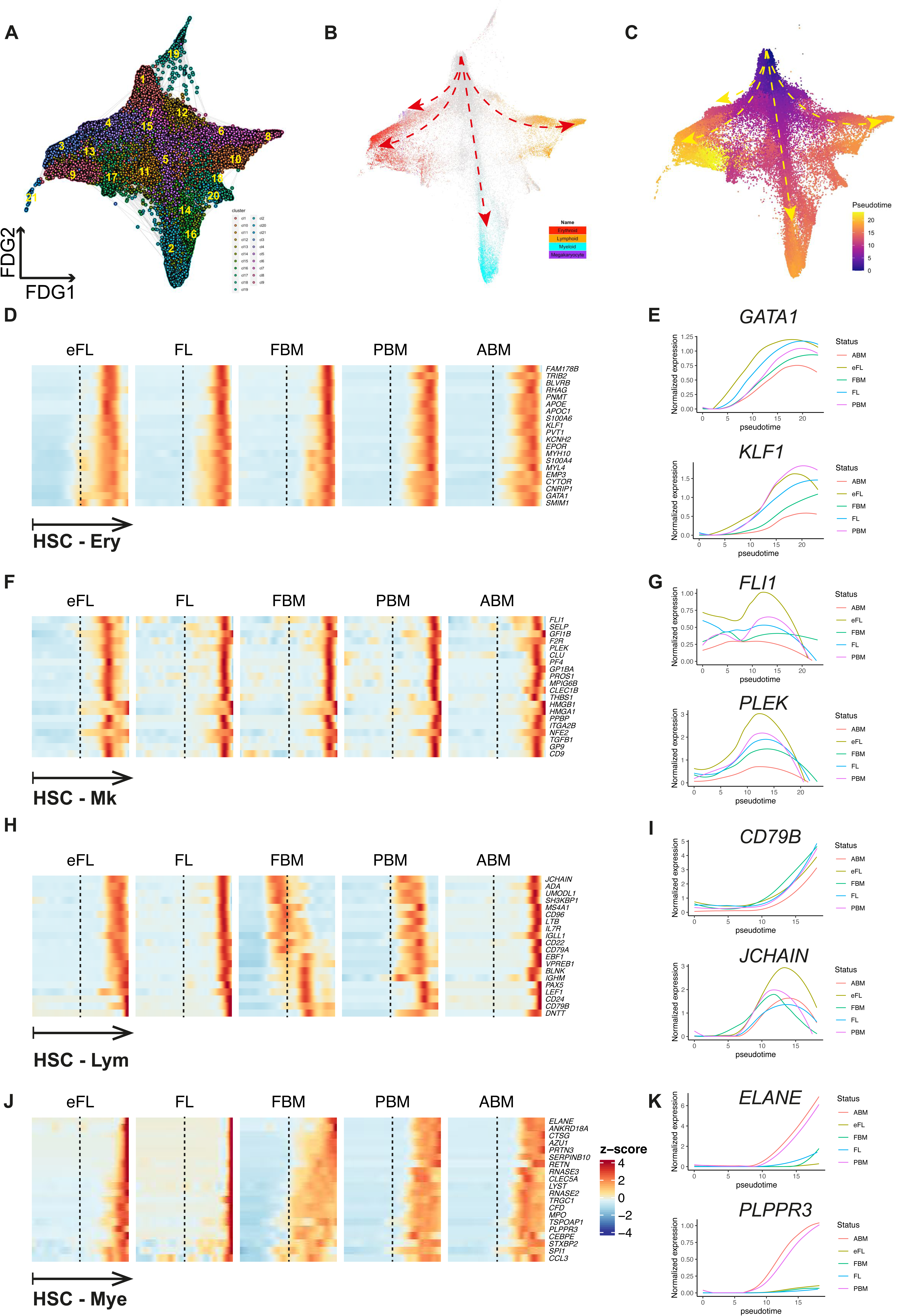
Four main differentiation trajectories were identified, and onset of lineage-affiliated transcriptional programs varied along ‘pseudotime’ between tissues. **(A-C)** Force-directed graph (FDG) showing **(A)** 21 Louvain clusters **(B)** superimposition of gene scores for 4 lineage gene sets **(C)** pseudotime score from Monocle3 analysis. Dashed arrows represent the four main trajectories of differentiation from HSC/MPP towards lymphoid, erythroid, megakaryocytic and myeloid progenitors. **(D, F, H, and J)** Expression of canonical lineage-affiliated genes along pseudotime from HSC to erythroid, megakaryocytic, lymphoid and myeloid trajectories for each tissue. **(E, G, I, and K)** Example erythroid, megakaryocytic, lymphoid and myeloid lineage-affiliated gene expressions along pseudotime for each tissue.

Monocle3 (Trapnell et al., 2014) analysis was performed to calculate pseudotime and identify differentiation paths for the four main trajectories (erythroid, megakaryocytic, lymphoid, and myeloid) (Suppl. Fig. 3D). To compare lineage differentiation trajectories across tissue types, the datasets were ‘down-sampled’ to control for differences in numbers of cells captured from each tissue, and cells were then ordered in pseudotime (Fig. 3D – 3K). The expression of erythroid and megakaryocyte-associated transcriptional program occurred earlier in pseudotime for eFL than other tissues (Fig. 3D – 3G). The expression of lymphoid-associated genes was markedly different between tissues across ontogeny, with onset of expression markedly earlier at the onset of bone marrow hematopoiesis in the 2^nd^ trimester fetus, and then was observed to occur progressively later in PBM and ABM cells (Fig. 3H and I). This included both early lymphoid genes and later B-cell specific genes, suggesting an accelerated B cell specification program in fetal and pediatric BM. Myeloid-associated gene expression occurred earlier in pseudotime for all BM tissues compared to eFL/FL and was particularly pronounced in FBM (Fig. 3J and K). The differences between matched FL and FBM from the same donors highlight differences in lineage specification between hematopoietic sites at the same developmental stage (Fig. 3D, F, H and J).

### Developmental stage drives global transcriptional differences between HSPCs, with enrichment of cell cycle genes in early fetal liver and inflammatory pathways in adult bone marrow

HSPCs from different developmental stages and hematopoietic sites showed distinct molecular profiles. A total of 4,094 genes was identified as differentially expressed (DE) upon pairwise comparison of the 5 tissue types, with the largest number of DE genes found between eFL and ABM (1,235 DE genes), and the smallest difference between FL and FBM samples from the same developmental stage (54 DE genes), suggesting that developmental stage is a more significant driver of global transcriptional differences than is the site of hematopoiesis (Suppl. Fig. 4A). Hallmark GSEA on differentially expressed genes between pairwise comparisons of selected tissues demonstrated enrichment of cell cycle related pathways (e.g. E2F target and G2M checkpoint) in eFL *vs*. FL, heme metabolism in FL *vs*. FBM, and multiple inflammatory response-related pathways in adult *vs*. paediatric BM (Suppl. Fig. 4B).

We also identified subsets of specific genes that were differentially expressed in each tissue when compared to all others (Suppl. Fig. 4C and Suppl. Table 5). Unsurprisingly, genes involved in controlling the cell cycle (*MYC)* and erythroid lineage-associated genes (*CNRIP1, TIMP3, and GATA2*) together with fetal-specific genes (*LIN28B*, and *IGF2BP3*) were highly expressed in eFL and FL (Copley et al., 2013; McWilliams et al., 2013; Yuan et al., 2012). *DHRS9*, a robust marker for regulatory or suppressive macrophages (Riquelme et al., 2017) was also specifically expressed in FL. FBM had strong expression of chemokine ligands (*CCL3, CCL3L1, CCL4*, and *CXCL8*) as well as B lymphoid genes (*VPREB3* and *CD83*) (O’Byrne et al., 2019) reflecting the strong myeloid/lymphoid skew in FBM. The postnatal samples PBM and ABM had relatively similar transcriptional profiles, with only 193 genes differentially expressed (Suppl. Fig. 4A). However, expression of *LAIR2*, a gene encoding an immunoglobulin superfamily receptor, was very specific for PBM perhaps associated with its role in the establishment of the innate immune system during childhood (Suppl. Fig 4C). Other genes expressed more highly in PBM vs. ABM included genes associated with early B-lymphoid development (*DNTT* and *FLT3*), the transcription factor and proto-oncogene (*ETV6*) that is frequently mutated in hematological malignancies, and genes involved in cell growth, proliferation and apoptosis (*YPEL3* and *AKR1C3*; Suppl. Fig. 4C). ABM strongly expressed kruppel like factors (*KLF3* and *KLF9)*, markers of DNA stress (*DDIT4*), a regulator of inflammation (*SOCS3*), the protein phosphatase *IER5* that regulates cell growth and stress resistance (Kawabata et al., 2015), and myeloid associated genes (*AREG* and *CEBPB*) compared to other tissues (Suppl. Fig. 4C). As expected, the switch from fetal to adult hemoglobin was also evident with the fetal gamma globin gene *HBG2* strongly expressed in FL in contrast to PBM and ABM where the beta globin gene (*HBB*) was the most prominently expressed beta globin cluster gene. A complete list of significantly differential genes across tissues is provided in Suppl. Table 5.

### Distinct gene regulatory networks in uncommitted HSPCs underlie differences in lineage specification between hematopoietic tissues

To examine the molecular regulators that underpin the differences in lineage-priming and cellular heterogeneity between hematopoietic tissues, we performed Single-Cell Regulatory Network Inference and Clustering (SCENIC) to evaluate the activity of gene regulatory networks (GRNs) (Aibar et al., 2017). This method enables identification of ‘regulons’, or genes that are co-expressed with transcription factors, with known direct binding targets based on *cis*-regulatory motif analysis. The activity score of each regulon was quantified in each cell using AUCell (Fig. 4A). This enabled identification of regulons that were specifically enriched in uncommitted HSC/MPP (ZNF467, HOXA9 and HOXB5); lymphoid progenitors (PAX5, TCF3, and TBX21); myeloid progenitors (SPI1, CEBPD, and RUNX1); erythroid progenitors specifically (HES6); megakaryocytic-erythroid progenitors (STAT5A and GATA1); eo/baso/mast cell progenitors (FEV and FOXD4L1); and megakaryocytic progenitors (MEF2C) (Fig. 4A).

**Figure 4.**
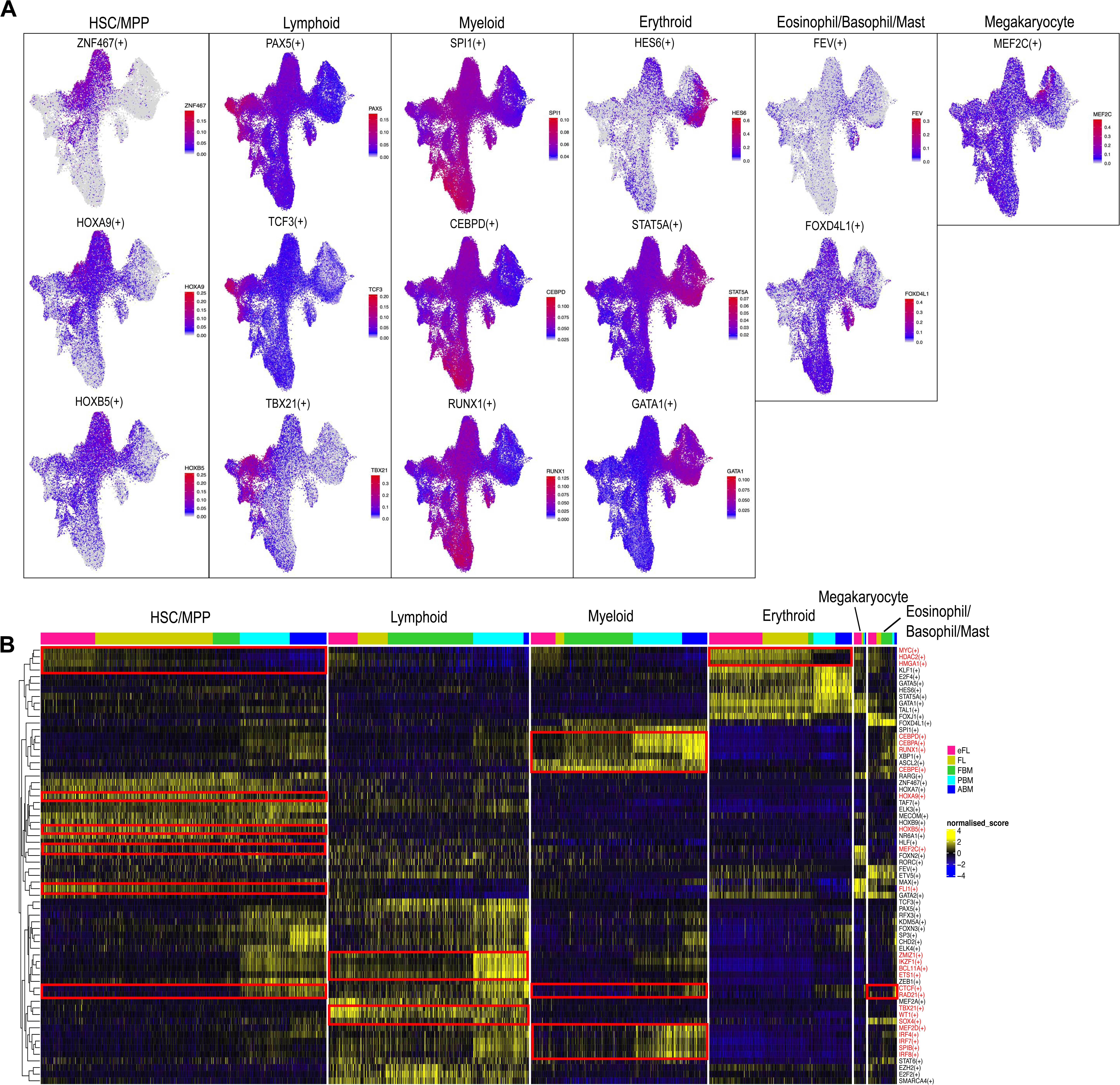
Distinct activity of gene regulatory networks (GRN) underlie differences in lineage specification across ontogeny. **(A)** UMAP plots superimposed with enrichment scores from SCENIC analysis for regulons specifically enriched in each lineage. HSC/MPP (ZNF467, HOXA9, HOXB5); lymphoid (PAX5, TCF3, TBX21); myeloid (SPI1, CEBPD, RUNX1); erythroid-specific (HES6); mega-erythroid (STAT5, GATA1); eo/baso/mast cell (FEV, FOXD4L); and megakaryocyte-specific (MEF2C). **(B)** Heatmap showing selected regulons that showed differential activity between lineages across tissues. The red boxes and names indicate regulons highlighted in the main manuscript.

The regulon activity was compared across tissue types for each HSPC subset (Fig. 4B). While the main differences in regulons were driven by the lineage specification of HSPC, there were tissue specific differences that were identifiable within uncommitted HSC/MPP and lineage-committed HSPC compartments. Uncommitted HSC/MPP from eFL had stronger enrichment of MYC, HDAC2, HMGA1, FLI1 and MEF2C, and expression of these regulons decreased substantially during development, suggesting a key role for these GRNs in promoting erythroid and megakaryocytic differentiation in eFL *vs*. hematopoietic tissues later in normal human development (Fig. 4B). Fetal HSC/MPP also showed enrichment for HOXA9 and HOXB5 regulons compared to postnatal tissues. There was an enrichment of lymphoid-associated GRNs in HSC/MPP from PBM, and this was accompanied by strong enrichment of ZMIZ1, IKZF1, BCL11A and ETS1 regulons in PBM lymphoid progenitors. TBX21, WT1 and SOX4 regulons were enriched in eFL lymphoid progenitors; MEF2D, SPIB, IRF4, 7 and 8, RUNX1 and CEBPA/D/E in postnatal myeloid progenitors; and MYC, HDAC2 and HMGA1 in fetal *vs*. postnatal erythroid progenitors. RAD21 and CTCF targets were specifically enriched in postnatal *vs*. prenatal HSC/MPP, erythroid and eo/baso/mast cell progenitors (Fig. 4B).

### Differences in the transcriptome of uncommitted HSPC through ontogeny

We next sought to investigate differences in the gene expression patterns of uncommitted HSPCs (HSC/MPP) through ontogeny, to identify possible cell-intrinsic factors underlying the changing differentiation bias during development. Cells from all tissues that were classified as HSC/MPP using an AUCell Score >0.15 were selected for further analysis (Fig. 5A and B). This demonstrated four distinct cell clusters – two minor subfractions that showed low-level enrichment of mega-erythroid and lympho-myeloid lineage gene signature sets, suggesting early lineage priming of some HSC/MPP (Fig. 5C), and, secondly, the non-lineage primed HSC/MPP were separated into two distinct subsets reflecting proliferative and quiescent cell states (Fig. 5D and E). After down sampling the data to analyse equal numbers of HSC/MPP from each tissue type, BM HSC/MPP from all developmental stages (FBM, PBM and ABM) showed myeloid-priming compared to FL samples, and PBM HSC/MPP were more lymphoid-primed than ABM, similar to our observations in total Lin-CD34+ HSPCs (Suppl. Fig. 5A).

**Figure 5.**
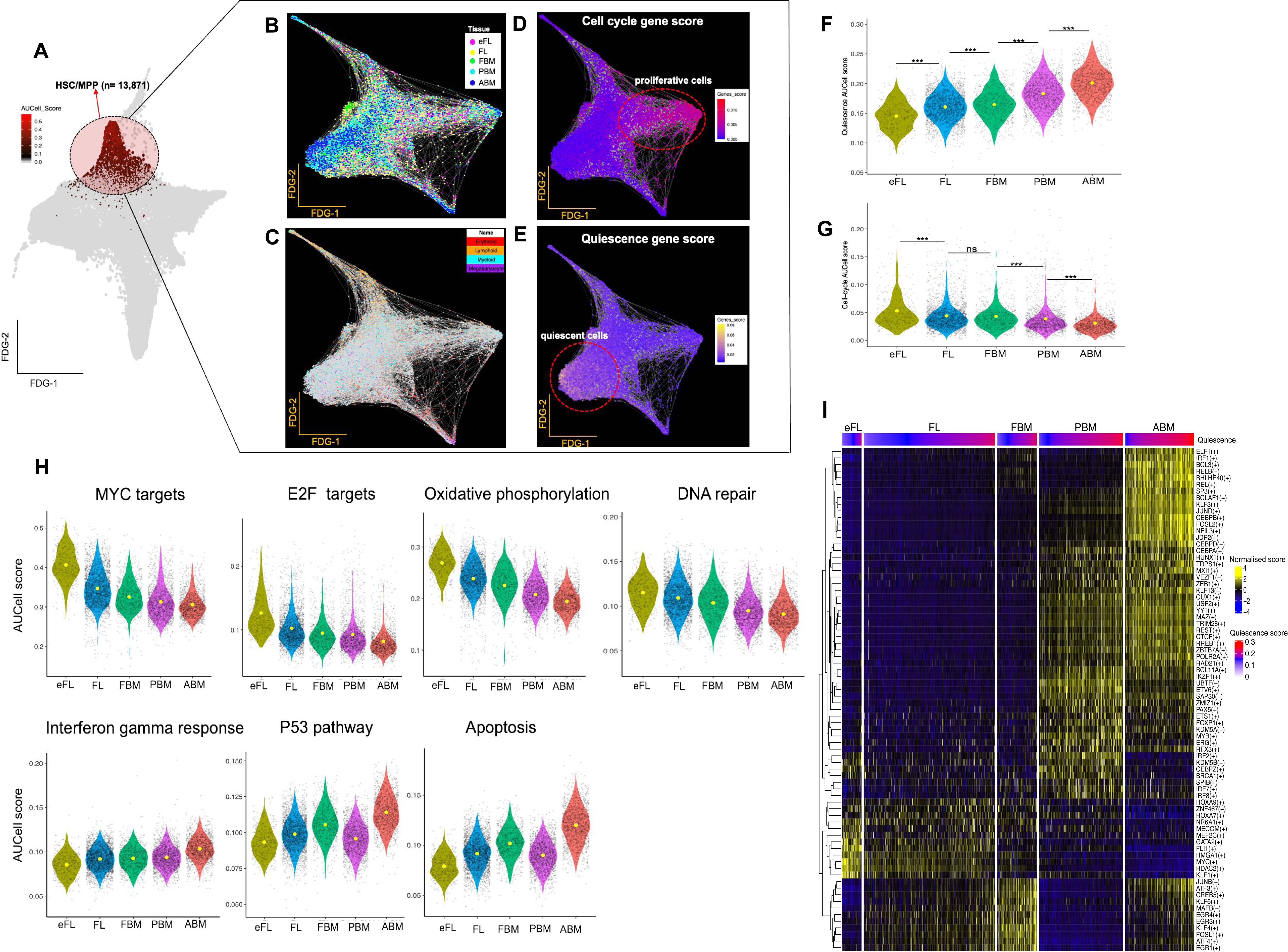
Uncommitted HSC/MPPs show early lineage priming and progression from cycling to quiescence over ontogeny, with differential GRN activity between tissues. **(A)** 13,871 cells were identified as HSC/MPP using an AUCell score > 0.15 for the HSC gene set) and visualised in a force-directed graph (FDG) with superimposition of **(B)** tissue of origin; **(C)** lineage gene set; **(D)** cell cycle gene expression scores and **(E)** quiescent gene expression scores. **(F and G)** AUCell scores for quiescence and cell cycle signature gene sets in a total of 6,398 HSC identified from different tissues. **(H)** Violin plots comparing AUCell scores from each selected HALLMARK gene set across tissues. **(I)** Differentially active regulons in HSC across tissues. Cells for each tissue were ranked by the quiescence score.

### Uncommitted HSCs show progression from proliferation to quiescence during ontogeny with enrichment of inflammatory signatures in adult bone marrow

To investigate the transcriptional profile of the most primitive HSC, cells with the very highest HSC AUCell score (> mean score) within the HSC/MPP compartment were taken forward for further characterisation (6,398 cells, Suppl. Fig. 5B). Quiescence and proliferation gene signatures (Suppl. Table 4) showed opposite changes through ontogeny, with ABM HSC showing the highest quiescence score and early FL HSC being most proliferative (Fig. 5F and G). Notably, this temporal pattern in cell cycling observed in HSCs over ontogeny was clearly distinct to that seen in the lineage-affiliated progenitor clusters (Suppl. Fig. 5C). The decrease in cell cycle score through ontogeny correlated with decreasing expression of MYC and E2F targets, oxidative phosphorylation, and DNA repair pathways. In contrast, increasing HSC quiescence correlated with inflammatory response, activation of P53 response and cell death associated pathways (Fig. 5H).

We next sought to determine if cell-intrinsic drivers of lineage priming and cell cycling were detectable in the most primitive HSCs. We therefore examined the regulons that were actively enriched in the 6,398 HSCs with the highest HSC AUCell score at different stages of ontogeny. eFL and FL HSC showed evidence of erythroid and megakaryocyte-specific regulons such as KLF1, GATA2, FLI1 and MEF2C (Fig. 5I). The FBM regulon was enriched for calcineurin-regulated NFAT-dependent transcription networks (FOSL1, JUNB, EGR3, EGR4 and MAFB) that are usually seen in lymphocytes, which might represent early lymphoid priming. PBM HSC showed a predominance of B lymphoid specific regulons, such as BCL11A, IKZF1, PAX5 and IRF8 compared to other tissues. The regulons specifically expressed in ABM HSCs were predominantly mediating inflammatory programs (e.g., interferon pathway (ELF1 and IRF1), NFKB and acute phase response (CEBPB), as well as genes previously reported as deregulated in aged HSCs (FOSL2 and JUND, Fig. 5I)(Lavrovsky et al., 2000; Schafer et al., 2018).

### Leveraging single cell transcriptomics to understand abnormal hematopoiesis

Finally, to demonstrate the utility of the dataset for understanding transcriptional perturbations in hematological diseases, we compared the normal ontogeny dataset to previously published single-cell datasets from two disease states. We chose datasets from hematological neoplasms that occur specifically in either early life (Juvenile Myelomonocytic Leukemia; JMML (Louka et al., 2021) or in adulthood (primary myelofibrosis; MF) (Psaila et al., 2020). We integrated the 57,489 Lin-CD34+ cells sampled from tissues across normal human ontogeny with 19,524 Lin-CD34+ cells from JMML patients (n=2) and 19,524 Lin-CD34+ cells from MF patients (n=15). The total dataset of 96,537 cells was then interrogated for lineage specification as for Fig. 2A. Compared to its normal counterpart (PBM), JMML HSPC had fewer lymphoid, erythroid and megakaryocytic progenitors (Fig. 6A) with majority of the HSPC being defined as HSC/MPP or myeloid progenitors (Fig. 6B). This is in keeping JMML being a myeloid neoplasm that originates in the earliest HSC compartment with an expansion of an abnormal CD38-CD90+ primitive HSPC population, with a reduced or abnormal erythroid, megakaryocytic and lymphoid output from JMML HSC *in vitro*, and a myeloid biased reconstitution *in vivo* (Louka et al., 2021). When compared to normal ABM, MF HSPCs showed dramatic expansion of megakaryocyte progenitors with almost complete absence of lymphoid progenitors (Fig. 6A and B), as previously reported (Psaila et al., 2020).

**Figure 6.**
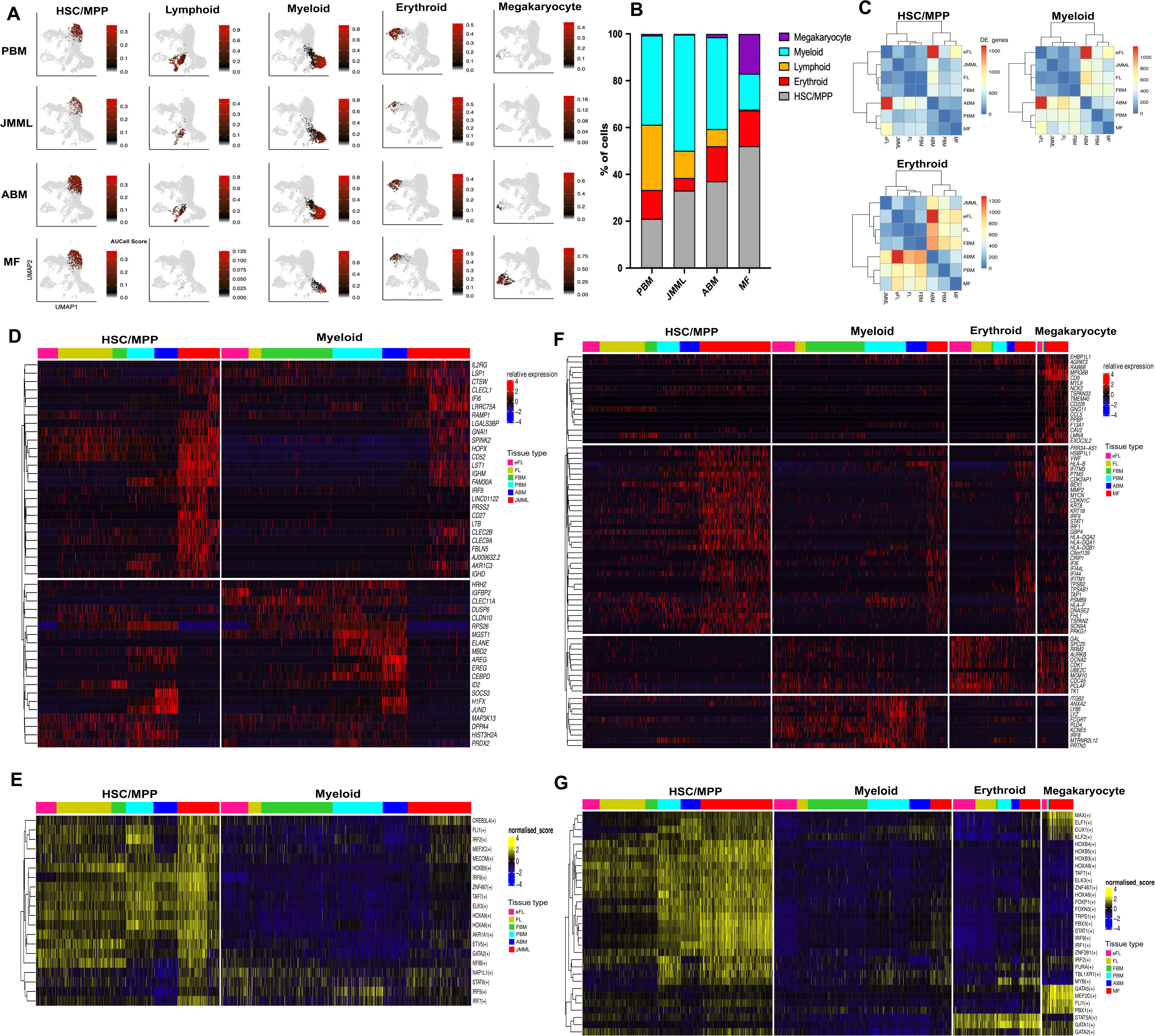
Concordance in gene expression between Juvenile Myelomonocytic Leukemia (JMML) HSPCs and normal fetal HSPCs; and increased inflammatory signaling seen in myelofibrosis (MF) and ABM in HSPCs. Data was derived from JMML (n=2, total cells=19,524) and MF (n=15, total cells=19,524) HSPC **(A)** UMAP plots showing lineage progenitors as identified by AUCell score >0.15 for ‘downsampled’ datasets to show 5,600 cells per tissue type. **(B)** The proportion of HSPCs (from all cells within each tissue), classified as HSC/MPP or lineage progenitors in JMML, MF and their normal counterparts. **(C)** Hierarchical clustering of the number of differentially expressed genes from pair-wise comparisons of tissues from normal ontogeny and JMML and MF for HSC/MPP, myeloid and erythroid compartments (adjusted *p*-value < 0.05, absolute log2FC > 1.5, and expressing cell frequency > 0.3 per tissue). **(D and E)** Top differentially expressed genes and regulons respectively, between JMML HSPCs and those from normal human ontogeny tissues for HSC/MPP and myeloid progenitors. **(F and G)** Comparison of top differentially expressed genes and regulons respectively, between MF HSPCs and those from normal human tissues through ontogeny for HSC/MPP and myeloid, erythroid and megakaryocyte lineage progenitors.

The developmentally-regulated molecular features of the progenitors in which blood cancers originate might drive the distinct biology of the disease at different ages. To understand whether the abnormal HSPC compartments in JMML and MF share features with normal counterparts from a particular developmental stage, we compared HSC/MPP and lineage-specific progenitors from disease states to their counterparts from all tissue types through ontogeny. Hierarchical clustering of the number of differential genes revealed that HSC/MPP, myeloid, and erythroid progenitors in childhood JMML clustered with fetal counterparts rather than postnatal PBM, whereas MF HSPC clustered with postnatal counterparts (Fig. 6C).

Differential gene expression analysis was performed to compare cells from JMML and MF to normal ontogeny in each lineage (Suppl. Table 6). JMML HSC/MPP showed higher expression of stem cell genes (e.g. *HOPX, SPINK2* and *CLEC9A*) compared to normal counterparts, and this ‘stemness’ signature was retained in JMML myeloid progenitors when compared to normal counterparts (Fig. 6D). Concomitantly, more mature myeloid genes such as *ELANE* and *CEBPD* were downregulated in the JMML myeloid progenitors compared to normal counterparts, suggesting a block in differentiation along the granulocytic/neutrophil lineage. Gene regulatory network analysis showed similarities of JMML HSC/MPP regulons with fetal HSC/MPP (FLI1, MEF2C, MECOM and GATA2), which were also enriched in JMML myeloid progenitors (Fig. 6E).

MF HSC/MPP showed high expression of immune and inflammatory pathways (*HLA* genes, *GBP4* and *IFITM1*), a matrix and collagen degrading enzyme (*MMP2*) and genes with oncogenic function (*MYCN*, *KRT8* and *KRT18*) that have been correlated with cancer progression and poor survival(Fortier et al., 2013; Lai et al., 2017). Megakaryocyte progenitors from MF showed markedly increased expression of a subset of MK-associated genes involved in cell-matrix interactions and cell adhesion (*CD9* and *MPIG6B)* when compared to the megakaryocyte progenitors from healthy control tissues. Similar to MF HSC/MPP, megakaryocyte progenitors in MF also showed increased expression of oncogenes (*RAB6B*), interferon inducible genes (*IFITM3*) and the cell cycle regulator *CDK2AP1* (Fig. 6F). Gene regulatory network analysis highlighted the activation of inflammatory signalling pathways (IRF9, IRF1, ELF1 and STAT1 regulons) and cell cycle (CUX1) in MF HSC/MPP and megakaryocyte progenitors, as well as resurgence of megakaryocyte-associated regulons (MEF2C, FLI1) that were most prominent in eFL HSC/MPP in the normal ontogeny dataset (Fig. 6G and Fig. 5I).

## DISCUSSION

Single cell approaches have been extensively applied to understand normal and perturbed hematopoiesis and to define human prenatal blood and immune cells (Park et al., 2020; Popescu et al., 2019; Ranzoni et al., 2021). These studies, together with other observations, have highlighted the predominance of erythroid lineage cells in human FL (Popescu et al., 2019) and B-lymphoid cells in FBM (O’Byrne et al., 2019; Popescu et al., 2019). Differences in the proliferative capacity of fetal and postnatal HSPC have been described using functional studies (Bowie et al., 2006; Copley et al., 2013; Lansdorp et al., 1993; Muench et al., 1994). A switch from multipotent to largely oligo/unipotent stem cells has also been documented between fetal and adult life (Notta et al., 2016). However, HSPCs sampled from over multiple timepoints over human ontogeny from 1^st^ trimester to adulthood at single cell resolution have not previously been reported.

In this study, we sought to define how hematopoiesis evolves during the human lifespan by interrogating a comprehensive transcriptomic dataset of HSPCs sampled from 5 hematopoietic tissues over 4 stages of ontogeny. Our data clearly demonstrate the changing frequencies of lineage-specific subsets that exist within the HSPC compartment. Notably, directly comparing FL and FBM HSPCs isolated from the same 2^nd^ trimester fetuses showed that while more than half of HSPCs in 2^nd^ trimester FL are uncommitted, >80% of HSPCs in the matched FBM are lineage primed, with dramatic expansion of myelo-lymphopoiesis in FBM. This enrichment of oligo/unipotent progenitors and switch in lineage output is likely to be driven by microenvironmental cues, as hematopoiesis migrates from FL to BM during the second trimester, although could also be explained by selective engraftment and subsequent expansion of lympho-myeloid progenitors in the BM microenvironment. We also found key differences in the gene regulatory pathways between tissues, supporting previous studies indicating that a complex interplay between cell intrinsic (Dykstra et al., 2011; Grover et al., 2016) and extrinsic factors (Ho et al., 2019) alter hematopoiesis and lineage specification during postnatal life. A marked and progressive increase in HSC quiescence was evident during postnatal development. As cell cycle may play a role in determining the fate of multipotent progenitors, at least for erythro-megakaryocytic lineage specification (Lu et al., 2018; Tusi et al., 2018), it is possible that this may also play an instructive role in the myeloid-biased hematopoiesis that is observed with increasing age.

Finally, mapping of two hematological malignancies onto normal hematopoiesis datasets allowed us to identify the similarities and differences in lineage specification and gene regulatory networks between these diseases and normal hematopoiesis through ontogeny. Such comparisons showed a persistence of fetal-like gene expression programs in the childhood disease JMML, supporting a fetal origin of this disease(Helsmoortel et al., 2016). Myelofibrosis, a cancer that typically presents in later adulthood, showed further exacerbation of inflammatory signalling pathways initiated in adult BM, together with altered cell-matrix interactions and activation of oncogenic programs, and resurgence of megakaryocyte-associated transcriptional signatures that were prominent in fetal liver but subsequently downregulated in later physiological development.

In conclusion, defining the key similarities and differences between hematopoietic tissues across normal human ontogeny has provided a key resource to study how the characteristics of the most primitive HSPC and their lineage fates change through the human lifetime. These results pave the way for a better understanding of hematopoiesis in normal human development, necessary to indicate the composition and likely long-term reconstitution ability of HSPC selected from donors of different ages for cell-based therapies, including gene editing and stem cell transplantation (Harrison et al., 1997; Holyoake et al., 1999; Hua et al., 2019; Muench et al., 1994; Nicolini et al., 1999), as well as the cellular and molecular underpinnings of age-specific vulnerabilities to the origin and evolution of certain disease states.

## ACKNOWLEDGEMENTS

We would like to acknowledge the contributions of the MRC WIMM Single Cell Facility and MRC-funded Oxford Consortium for Single-Cell Biology (MR/M00919X/1); WIMM Flow Cytometry Facility which is supported by the MRC HIU, MRC MHU (MC_UU_12009), NIHR Oxford BRC, Kay Kendall Leukemia Fund (KKL1057), John Fell Fund (131/030 and 101/517), the EPA fund (CF182 and CF170) and by the WIMM Strategic Alliance awards G0902418 and MC_UU_12025. We thank the High-Throughput Genomics Group at the Wellcome Trust Centre for Human Genetics (funded by Wellcome Trust Grant Reference 090532/Z/ 09/Z) and the NIH Intramural Sequencing Core and the National institutes of Health Research (NIHR) Oxford Biomedical Research Centre (BRC). The human fetal material was provided by the Joint MRC/Wellcome Trust Grant 099175/Z/ 12/Z Human Developmental Biology Resource (http://hdbr.org). We also thank the donors who kindly provided samples for this research. A.R. was supported by a Bloodwise Clinician Scientist Fellowship (grants: 14041 and 17001), Wellcome Trust Clinical Research Career Development Fellowship (216632/Z/19/Z), MRC Discovery award (MRCDA 0816-11), Lady Tata Memorial International Fellowship, and EHA-ASH Translational Research Training in Hematology Fellowship. B.P. was supported by a Wellcome Clinical Research Career Development Fellowship, a LAB282 award, an Oxford BRC Senior Research Fellowship and a Cancer Research UK Advanced Clinician Scientist Fellowship. S.T. was supported by Oxford-Bristol Myers Squibb (BMS) Fellowship.

## AUTHOR CONTRIBUTIONS

A.R, D.I., S.O’B, N. E., J.O’S, G.B, P. H. provided samples and/or performed experiments. G.W., E. H., W. X. W., A.R.-M. designed and performed bioinformatic analyses. A. K., A.J.M, D.B. and I.R. contributed to project oversight. A.R., B.P and S.T. conceptualised the project, analysed and interpreted data and wrote the manuscript. S.T. designed and developed the SingCellaR software. All authors read and approved the manuscript.

## METHODS

### Human samples

Donated fetal tissue (1^st^ trimester FL and matched 2^nd^ trimester FL and FBM) was provided by the Human Developmental Biology Resource (HDBR, www.hdbr.org). Fetal tissues were transported to the laboratory at 4°C and processed immediately as described previously (O’Byrne et al., 2019; Roy et al., 2012). Adult BM mononuclear cells were purchased from StemCell Technologies, Canada (cat no. 17001). Normal pediatric bone marrow was prospectively collected in accordance with the Declaration of Helsinki for sample collection and use in research. After filtering through a 70 micron cell strainer, samples were red cell and granulocyte depleted by density gradient separation using Ficoll-Paque PLUS (GE Healthcare Life Sciences, cat. no. 17-5442-03) and CD34 enrichment was carried out on freshly isolated mononuclear cells (MNC) from some of the samples using a Miltenyi CD34 MicroBead kit and MACS system (Miltenyi Biotech, cat. no. 130-046-703). Developmental stage (age) and sex can be found in Suppl. Table 1.

### Fluorescent activated cell sorting (FACS) staining, analysis and cell isolation

Cells were stained with fluorophore-conjugated monoclonal antibodies (mAb; see Suppl. Table 7) in PBS with 2% FBS and 1mM EDTA for 30 minutes followed by two washes. FACS-sorting was performed using a Becton Dickinson Aria III or Fusion 2 as previously described (Psaila et al., 2020).

### High-throughput single-cell RNA-sequencing (10x Chromium)

15,000-24,000 Lin-CD34+ cells were FACS sorted from each sample, and processed as described in (Psaila et al., 2020).

### 10x Genomics single-cell RNA sequencing

Demultiplexed FASTQ files were aligned to the human reference genome (GRCh38/hg38) using Cell Ranger software (version 3.0.1) from 10x Genomics. The Cell Ranger ‘‘count’’ standard pipeline was used to obtain the expression matrix of Unique Molecular Identifier (UMI) for each individual sample.

### Data and code availability

All raw and processed sequencing data generated in this study have been submitted to the NCBI Gene Expression Omnibus (GEO; https://www.ncbi.nlm.nih.gov/geo/). The data will be released after the official publishing of this study. The JMML and MF datasets were previously published and are available under accession numbers: GSE111895 and GSE144568 respectively. SingCellaR open-source codes are available and maintained on GitHub https://github.com/supatt-lab/SingCellaR.

### Data processing and filtering of HSPC dataset

We used SingCellaR (https://github.com/supatt-lab/SingCellaR) to process each sample individually. The function ‘load_matrices_from_cellranger’ was used to read in data matrices from the Cell Ranger output. Cell and gene filtering was performed by assessing QC plots using the ‘plot_cells_annotation’ function. Cells meeting the following QC parameters were included in analyses (Suppl. Table S1): UMI counts > 1,000 and ≤ maximum UMIs; number of detected genes > 500 and ≤ maximum number of detected genes; the percentage of mitochondrial gene expression ≤ limited percentage of mitochondrial gene expression (10% or 20% depending on an individual sample). Genes expressed in at least 10 cells were included. After filtering according to these criteria, 57,489 cells passed quality control (Suppl. Table 1) and were included in downstream analyses.

### Data integration

To perform data integration, the ‘SingCellaR_int’ R object was created. R object file names from individual samples were required as the input for the object. The function ‘preprocess_integration’ was performed to combine all of UMIs from all samples and cluster together with marker gene information into a single integrated R object. Using the function ‘get_variable_genes_by_fitting_GLM_model’ as previously described (Psaila et al., 2020), 949 highly variable genes, after removing ribosomal and mitochondrial genes, were identified and used for PCA analysis of the integrated dataset. Data normalization (using scaled UMI counts by normalizing each library size to 10,000 and transforming to the log scale) and dimensional reduction were performed using the function ‘normalize_UMI’ and ‘runPCA’, with the fast PCA analysis from the IRLBA package. To perform integration, the function ‘runSupervised_Harmony’ (implemented in SingCellaR) was run by using 40 principal components (PCs) as determined by the PCA elbow plot generated using the ‘plot_PCA_Elbowplot’ function. runSupervised_Harmony function automatically groups marker genes for each cluster per sample and builds up a gene set database for GSEA analysis. An iterative GSEA is then performed to test significance level and to calculate positive enrichment scores for marker gene sets for each cluster by querying the marker genes identified per cluster against the whole gene set database. A matrix of cluster matching scores from GSEA enrichment values is generated and used as the input for an automated hierarchical clustering with the ‘cutree’ function. This can be used to estimate the number of likely matched and unmatched clusters from all samples. Common and unique clusters across samples are then identified from the hierarchical clustering tree using the ‘cutree’ function. A prior probability matrix is calculated for each single cell belonging to the common and unique clusters. Finally, the probability matrix is provided as the input ‘cluster_prior’ together with the ‘donor’ and ‘batch’ as the covariates for the subsequent Harmony analysis (Korsunsky et al., 2019).

### Benchmarking distinct integrative methods for HSPC dataset

We performed kBET and LISI methods (Buttner et al., 2019; Korsunsky et al., 2019) on our HSPC dataset (Suppl. Fig. 1) to benchmark the SingCellaR integrative method against previously published methods for integrating datasets. To run kBET, we annotated clusters of HSPCs using AUCell score (Aibar et al., 2017). We selected cells with a high AUCell score (> 0.15) of HSC/MPP, myeloid, lymphoid, erythroid, megakaryocyte, eosinophil/basophil/mast and endothelial progenitor cell gene signatures (Suppl. Table 4) and classified them into 7 groups. Due to strong AUC cell score, indicating strong expression of signature genes per each selected lineage progenitor, we would expect that each group of cells should be aggregated well together when applied integrative methods. We next performed each data integration method and UMAP analysis. UMAP 2D-coordinates for all methods were used as the input matrices for kBET analysis. kBET analysis was run for each group of annotated lineage progenitor cells by subsampling 1,000 cells/group and donor information was used as the batch of interest. kBET average acceptance rate per group of HSPCs was calculated and plotted for each integration method (Suppl. Fig. 1D). For LISI, the function “compute_lisi” was performed with the input UMAP 2D-coordinates together with donor and annotated lineage information. iLISI and cLISI scores were calculated and plotted for each integration method (Suppl. Fig. 1E).

### Data visualisation with lineage signature gene sets

After data integration, data embedding methods were performed for visualisation using SingCellaR. These included UMAP, force-directed graph, and diffusion map using functions ‘runUMAP’, ‘runFA2_ForceDirectedGraph’, and ‘runDiffusionMap’. We investigated the expression of lineage signature gene sets superimposed on top of those embeddings. Lineage signature gene sets were collated by curating known canonical lineage markers selected from multiple published hematopoiesis datasets (Suppl. Table 3).

SingCellaR calculates a lineage gene score for each cell based on the average gene expression of each gene set. Represented colours for gene sets can be assigned automatically or by user-defined colours. Transparency factors for selected colours are calculated from normalised expression values across cell types. SingCellaR uses ‘ggplot2’ functionality for adding the dynamic alpha parameter values to ‘‘geom_point’’ to control the transparency of colours.

### Clustering analysis, marker gene identification, and cell type annotation

Clustering analysis was performed using the function ‘identifyClusters’ in SingCellaR with the integrative embeddings as the input together with the ‘cosine’ as a distant metric and local k-nearest neighbour (KNN) equal to 30. SingCellaR clusters cells using k-nearest neighbour approach implemented by a fast KNN algorithm from ‘RcppAnnoy’ package. After nearest neighbours are identified, the weighted graph is created with weight values calculated from normalised shared number of the nearest neighbours. The ‘louvain’ community detection method implemented by igraph package was applied to identify clusters. To identify genes differentially expressed in each cluster, the function ‘findMarkerGenes’ was performed. SingCellaR uses a standard nonparametric Wilcoxon test on log-transformed, normalised UMIs to compare expression level. Fisher’s exact test was used to compare the expressing cell frequency of each gene as previously described (Giustacchini et al., 2017). *P*-values generated from both tests were then combined using Fisher’s method and adjusted using the Benjamini-Hochberg (BH) correction. Genes expressed by each individual cluster are compared to all other clusters and differential genes defined as an absolute log2 fold change of ≥ 1.5 and adjusted *P-*value of < 0.05, with the fraction of expressing cell frequency of > 0.3. Differentially expressed genes were ranked using *P*-values and log2FC to select the top differential genes per cluster.

To annotate clusters, the function ‘identifyGSEAPrerankedGenes’ was used to pre-rank genes obtained from differential gene expression analysis comparing each individual cluster with all other clusters. Gene ranking scores were calculated as the log2 of expression fold-change multiplied by −log10 of the adjusted p-value. To run GSEA, the function ‘Run_fGSEA_for_multiple_comparisons’ was performed using the fGSEA package (Korotkevich et al., 2019). The function ‘plot_heatmap_for_fGSEA_all_clusters’ was used to perform hierarchical clustering and to visualise GSEA positive enrichment scores from all clusters on the heatmap. This cell annotation heatmap (Fig. 1C) together with the identified top marker genes (Suppl. Table 2) and manual curation was used to identify cell clusters.

### Gene set enrichment analysis

SingCellaR provides functions for GSEA analysis. These functions include: ‘Run_fGSEA_analysis’ used to compare two groups of cells; ‘Run_fGSEA_for_a_selected_cluster_vs_the_rest_of_clusters’ to compare any selected cluster against all other clusters; ‘Run_fGSEA_for_multiple_comparisons’ function to performing GSEA on multiple comparisons. Curated lineage signature gene sets used here are listed in Suppl. Table 3. The HALLMARK gene set was downloaded from MSigDB (https://www.broadinstitute.org/gsea/msigdb/collections.jsp). Genes were pre-ranked using the function ‘identifyGSEAPrerankedGenes’.

### Lineage progenitor quantification

AUCell (Aibar et al., 2017) was used to quantify the number of cells affiliated to each lineage (HSC/MPP, myeloid, lymphoid, erythroid, megakaryocyte, and eosinophil/basophil/mast) among all HSPCs). To run AUCell, the function ‘Build_AUCell_Rankings’ was performed followed by ‘Run_AUCell’ with the gene matrix transposed (GMT) file of lineage gene sets. We manually inspected different AUCell thresholds per each lineage. We selected cells with AUCell score > 0.15 for each lineage and calculated the proportion of cells in each lineage progenitor subset and between tissues.

### Pseudotime trajectory analysis

We used Monocle3 (version: 0.2.3.0) combining with the SingCellaR results to identify trajectories of the entire dataset. Raw UMI count data and clustering annotations were extracted from SingCellaR object to build a Monocle ‘cds’ object. We first used ‘preprocess_cds’ function to normalise the data. The UMAP and forced-directed graph dimensional reduction and clustering results slots were obtained from SingCellaR analyses. The trajectory was identified using “learn_graph” function on the UMAP reduction embeddings, representing paths along different lineages. “Order_cells” function was used to calculate pseudotime and the root node was defined using function “get_earliest_principal_node” on the UMAP reduction embeddings. The trajectory was defined from the graph plot as: 1) HSC – Ery: cl1-cl4-cl3-cl9; 2) HSC – Lym: cl1-cl7-cl6-cl8; 3) HSC – MKE: cl1-cl4-cl13; 4) HSC – Mye: cl1-cl7-cl5-cl2. We further downsampled the number of cells to control for differences in numbers of cells captured from each tissue (HSC – Ery: 1,650 cells per tissue; HSC – Lym: 1,781 cells per tissue; HSC – MKE: 1,363 cells per tissue; and HSC – Mye: 2,715 cells per tissue). Selected marker genes for each lineage were presented in heatmaps using the ComplexHeatmap package and line plot using ggplot2.

### Single-cell regulatory network inference and clustering (SCENIC) analysis

We used pyscenic (version 0.10.4) to perform single-cell regulatory network analysis. We performed the analysis by following the protocol steps described in SCENIC workflow (Van de Sande et al., 2020). We analysed the cells corresponding to uncommitted HSC/MPP and main lineage progenitor sub-types described as the AUCell score of > 0.15 (Fig. 2A). We first run the python script ‘arboreto_with_multiprocessing.py’ using the ‘grnboost2’ method followed by running the ‘pyscenic’ using default parameters with the database file ‘hg38 refseq-r80 10kb_up_and_down_tss.mc9nr.feather’ and the motif information file ‘motifs-v9-nr.hgnc-m0.001-o0.0.tbl’. The AUCell analysis was further performed using ‘pyscenic aucell’ function with parameters ‘rank_threshold’ 5000, ‘auc_threshold’ 0.05 and ‘nes_threshold’ 3. Identified regulons from pyscenic were further selected based on the average AUC score across cells > 0.02 and the number of genes in each regulon > 10. Differential regulons were selected using the visual inspection on the heatmap plot of normalised AUCell scores across tissues or lineages.

## SUPPLEMENTAL INFORMATION

**Supplemental figures and legends (Suppl. Figs. 1-5).** Compiled supplementary figures and legends referenced in the main text.

**Supplemental Tables 1-4.** Supplemental Table 1 – sample information and cell QC; Supplemental Table 2 – top 30 differentially expressed genes per cluster; Supplemental Table 3 – gene-sets from previous studies relevant to hematopoiesis; and Supplemental Table 4 – gene sets mainly used in lineage scoring and AUC cell analyses in this study.

**Supplemental Table 5.** A complete list of significantly differential genes across tissues.

**Supplemental Table 6.** A complete list of differentially expressed genes between JMML and MF versus normal ontogeny in each lineage.

**Supplemental Table 7.** A list of antibodies used for flow cytometry.

## Supplemental Information

### Supplemental Figure Legends

**Supplemental Figure 1.**
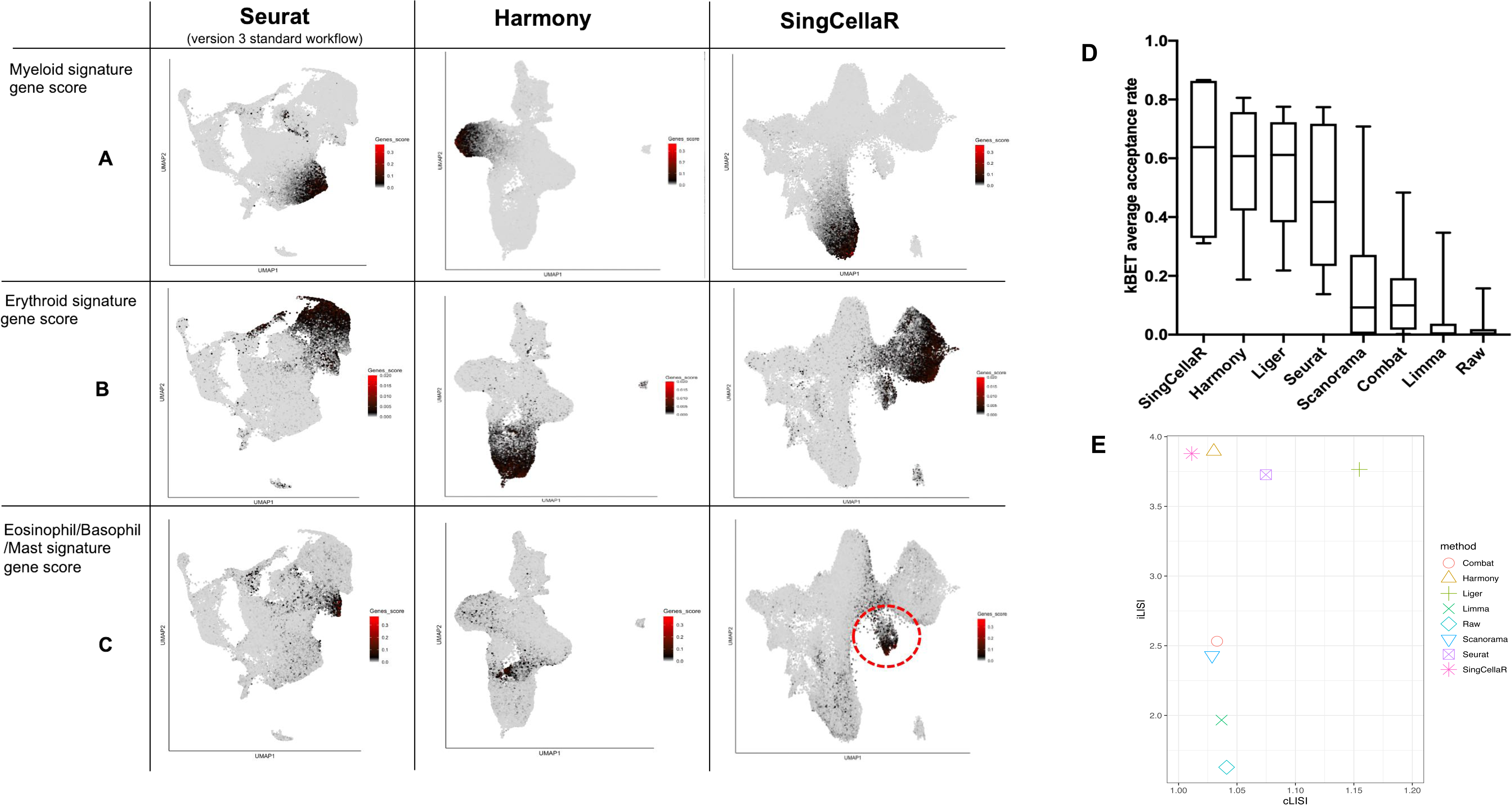
Optimal resolution of lineage progenitor clusters was obtained using a ‘supervised Harmony’ integration approach, implemented in SingCellaR. UMAP plots of selected integration methods (Seurat, Harmony and SingCellaR) that performed well for this HSPC dataset. Lineage signature gene scores for **(A)** myeloid, **(B)**, erythroid and **(C)** eosinophil/basophil/mast progenitors are superimposed on the UMAP plots, with a dashed circle highlighting the cluster of eosinophil/basophil/mast progenitor cells that is resolved only using SingCellaR. **(D and E)** Objective measures of integration for each method **(D)** Boxplot of kBET average acceptance rate score and **(E)** iLISI and cLISI scores. X-axis represents cLISI score. Y-axis represents iLISI score. A higher kBET and iLISI score indicates better data integration, and accurate integration should result in a cLISI score of 1.

**Supplemental Figure 2.**
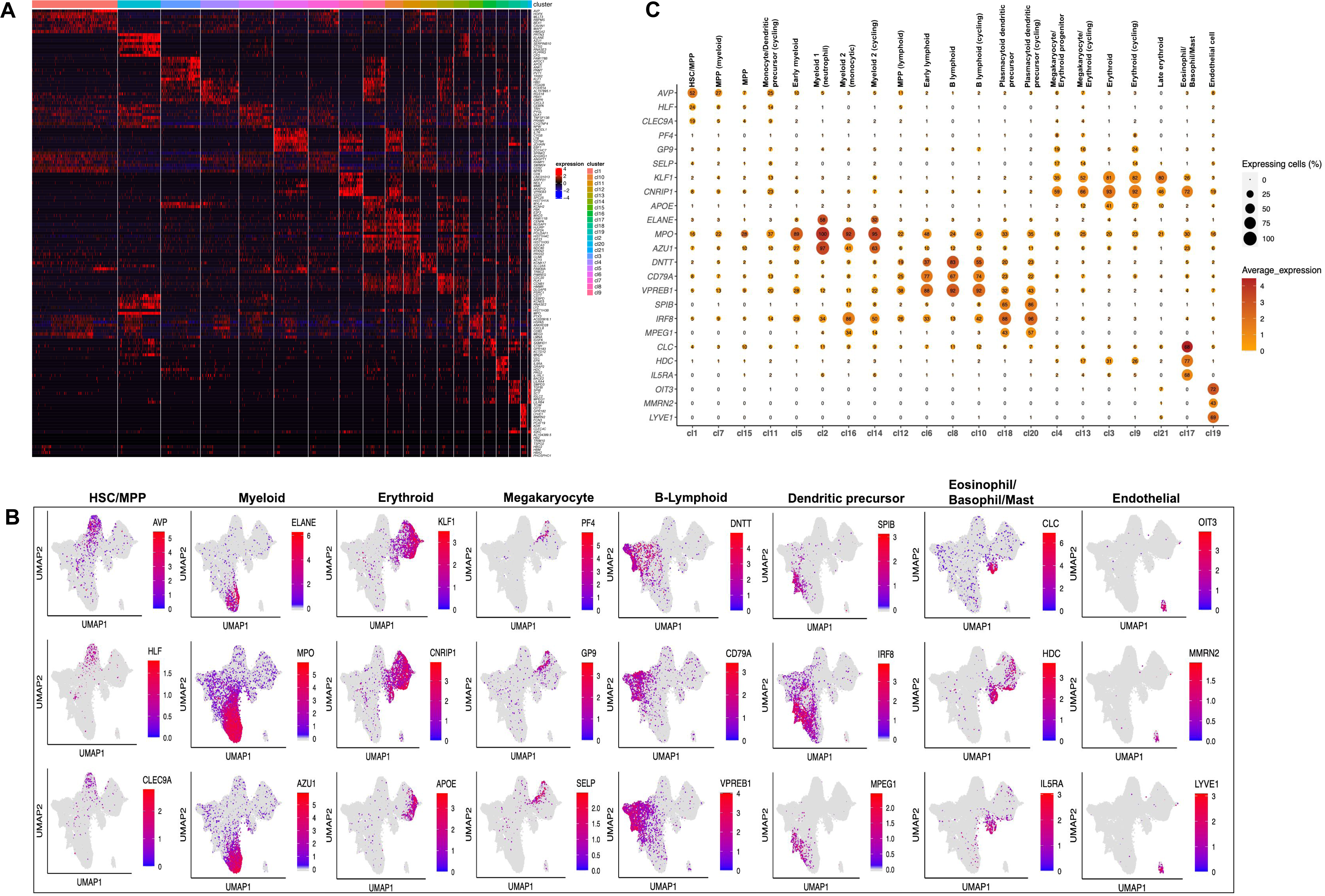
Identification of cell clusters using differentially expressed genes. **(A)** Heatmap showing relative expression of the top 8 differentially expressed genes for each cluster. Cluster IDs are ranked from left (Cluster 1) to right (Cluster 21). **(B)** UMAP plots displaying the expression of canonical lineage marker genes. **(C)** Bubble plots showing the expression of lineage marker genes for each cluster. The size of the dot represents the percentage of expressing cell frequency. The number inside the dot is the percentage of expressing cells.

**Supplemental Figure 3.**
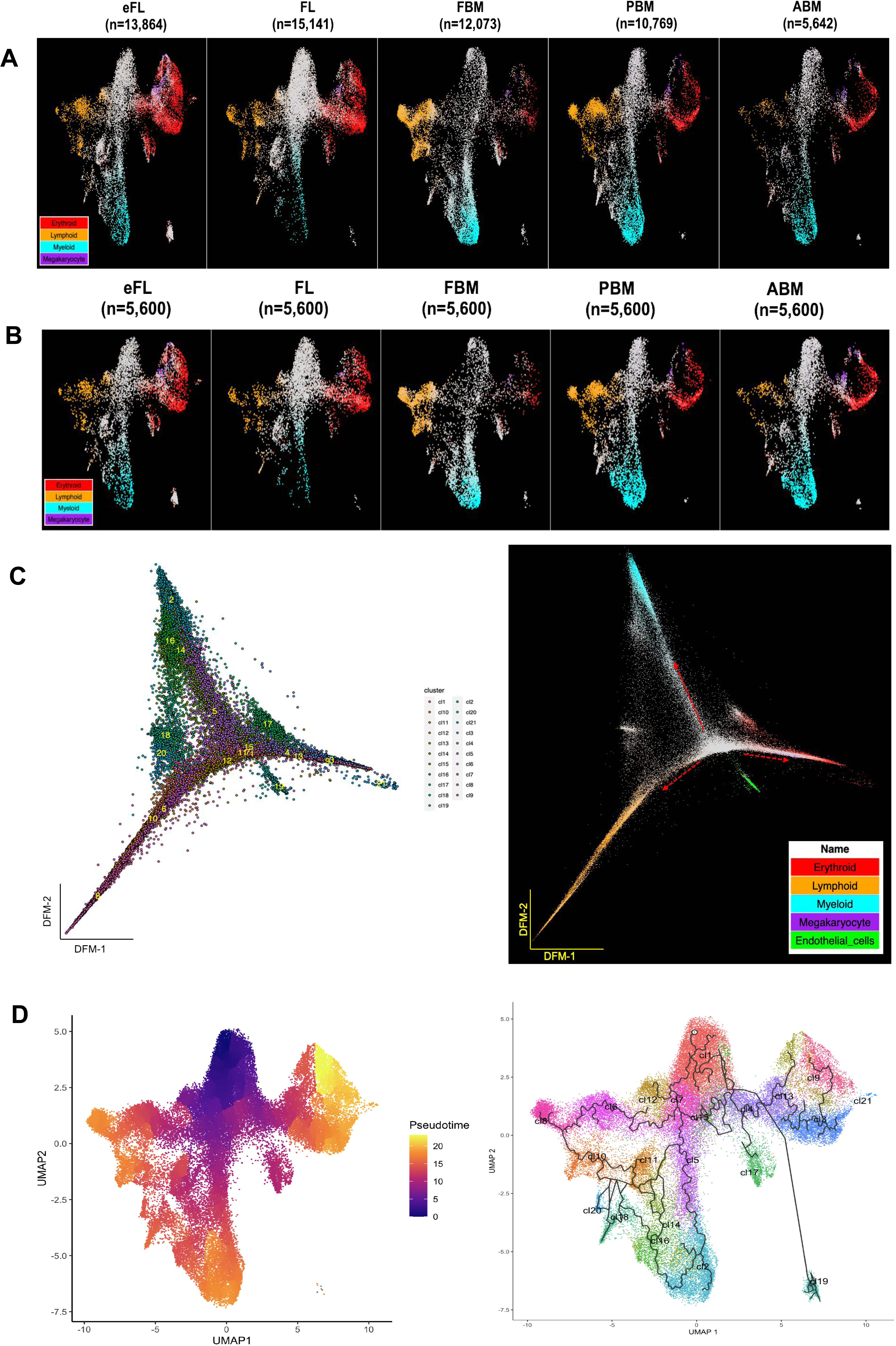
Dynamic changes in the cellular composition of HSPCs across tissues and differentiation trajectory analyses. UMAP plots overlaid with lineage gene signature scores for HSC/MPP (grey); erythroid (red); lymphoid (yellow); myeloid (cyan) and megakaryocyte (purple) **(A)** showing all cells in each tissue **(B)** showing down-sampled cells (5600 cells per tissue). **(C)** HSPCs differentiation trajectories on a diffusion map. Left panel, diffusion map with cluster ID information. Right panel, diffusion map with the superimposition of 5 lineages progenitor gene sets. Red - erythroid; yellow - lymphoid; cyan - myeloid; purple - megakaryocyte; green - endothelial cells. The arrows indicate the main trajectories of cell differentiation from HSC/MPP towards lymphoid, erythroid/megakaryocyte, and myeloid differentiation. **(D)** Pseudotime and trajectory graph using Monocle3 analysis. Left panel, UMAP-plot superimposed with pseudotime scale. Dark-blue indicates the starting state and yellow or red indicates the terminal state of the cellular trajectories. Right panel, UMAP-plot with clusters and identified trajectory paths based on the monocle analysis.

**Supplemental Figure 4.**
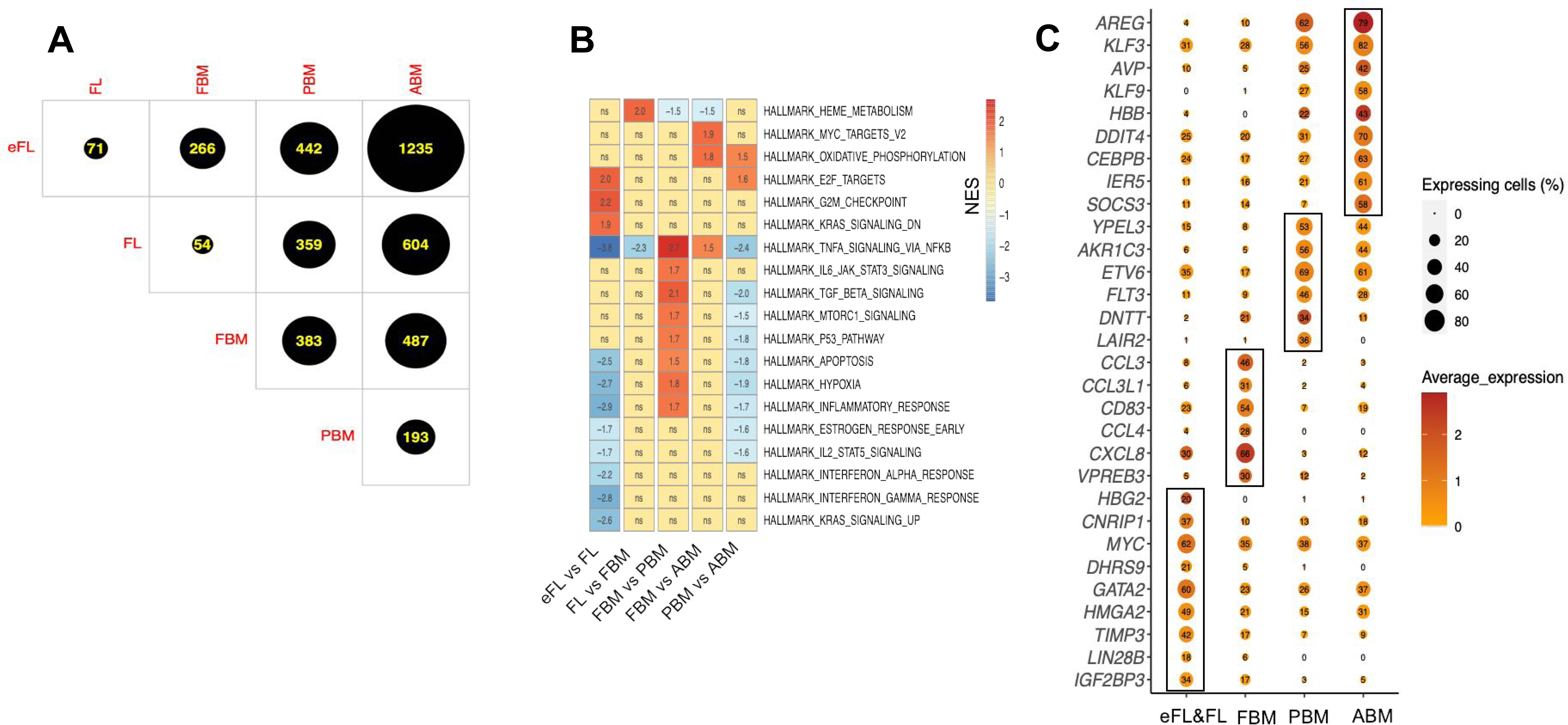
Dynamic changes in HSPC gene expression programs across human ontogeny and the trajectory analysis of selected HSC/MPPs across tissues. **(A)** The number of differentially expressed genes between tissues in pair-wise comparisons. **(B)** Heatmap of gene set enrichment scores for HALLMARK gene sets for the comparison of 5 selected pairs of tissues – eFL *vs*. FL, FL *vs*. FBM, FBM *vs*. PBM, FBM *vs*. ABM, and PBM *vs*. ABM. The gradient of colors represents the normalised enrichment score (NES) provided by GSEA. Red – gene sets enriched in the first tissue of the pair (positive values); blue – enrichment score for the second tissue of the pair (negative values). Yellow – no significant difference (‘ns’). **(C)** Selected tissue-specific genes that are highly expressed in each tissue.

**Supplemental Figure 5.**
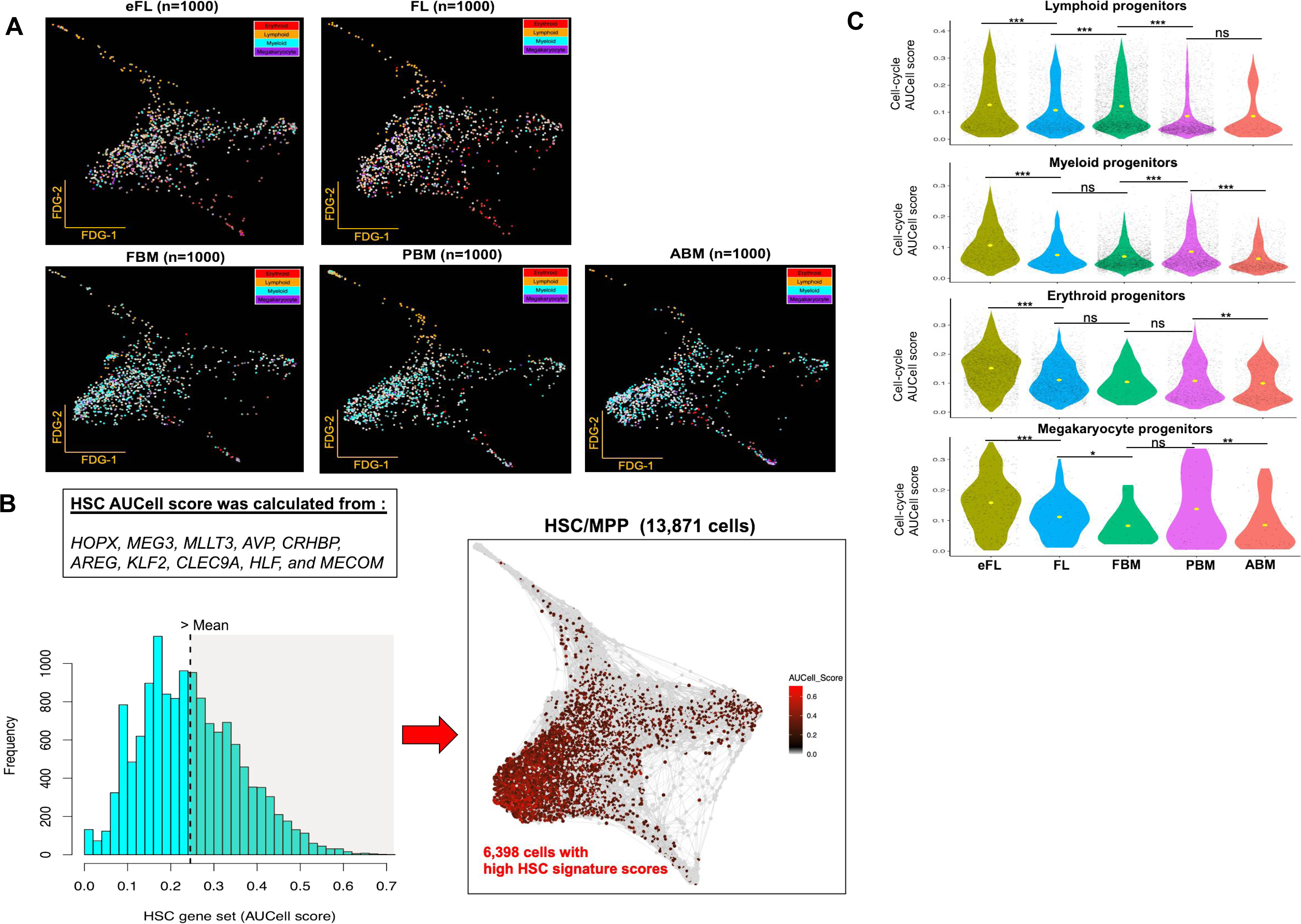
Selection of primitive HSCs and comparison of AUCell scores for selected gene sets across tissues. **(A)** Force-directed graph trajectory analysis of selected HSC/MPPs across tissues following ‘down-sampling’ to 1,000 cells per tissue. Lineage signature gene expression is superimposed on top of the graph. Red - erythroid; yellow - lymphoid; cyan -myeloid; purple – megakaryocyte progenitors. **(B)** Left panel, the AUCell score distribution calculated from 10 curated HSC genes. 6,398 cells had higher HSC score than the average (>2.5) and were selected for downstream analyses. Right panel, force-directed graph represents selected HSC (black and red cells) in the trajectory of HSC/MPP compartment. **(C)** Violin plots comparing cell cycle AUCell scores in different lineages across tissues. Significance level shown as obtained using a Wilcoxon test (* = *P* ≤ 0.05; ** = *P* ≤ 0.01; *** = *P* ≤ 0.001; and ‘ns’ = not significant).

